# The human cytomegalovirus G-protein coupled receptor US28 promotes latency by attenuating c-fos

**DOI:** 10.1101/434605

**Authors:** Benjamin A. Krishna, Monica S. Humby, William E. Miller, Christine M. O’Connor

**Author notes:** 9500 Euclid Ave, Mail Stop NE50, Lerner Research Institute, Cleveland Clinic, Cleveland, OH 44016.

## Abstract

Human cytomegalovirus (HCMV) is a ubiquitous pathogen that undergoes latency in cells of the hematopoietic compartment, though the mechanisms underlying establishment and maintenance of latency remain elusive. We previously reported that the HCMV-encoded G-protein coupled receptor (GPCR) homolog, *US28* is required for successful latent infection. We now show that US28 protein (pUS28) provided *in trans* complements the US28Δ lytic phenotype in myeloid cells, suggesting that sustained US28 expression is necessary for long-term latency. Furthermore, expression of pUS28 at the time of infection represses transcription from the major immediate early promoter (MIEP) within 24 hours. However, this repression is only maintained in the presence of continual pUS28 expression provided *in trans*. Our data also reveal that pUS28-mediated signaling attenuates both expression and phosphorylation of cellular fos (c-fos), an AP-1 transcription factor subunit, to repress MIEP-driven transcription. AP-1 binds to the MIEP and promotes lytic replication, and in line with this, we find that US28Δ infection results in an increase in AP-1 binding to the MIEP, compared to wild type latent infection. Pharmacological inhibition of c-fos represses the MIEP during US28Δ infection to similar levels we observe during wild type latent infection. Together, our data reveal that US28 is required for both establishment and long-term maintenance of HCMV latency, which is modulated, at least in part, by repressing functional AP-1 binding to the MIEP.

**Significance Statement:** Human cytomegalovirus (HCMV) is a wise-spread pathogen that remains with an individual for life in a quiescent/latent state, posing little threat to an otherwise healthy person. However, when an individual’s immune system is severely compromised, HCMV can reactivate to its active/lytic state, resulting in viral spread and disease that is often fatal. The biological mechanisms underlying HCMV latency and reactivation remain poorly understood. Herein we show that the viral-encoded G-protein coupled receptor (GPCR) *US28* aids in the establishment and the maintenance of viral latency. Furthermore, we find that US28 modulates host cell proteins to suppress viral processes associated with active/lytic replication, thereby promoting latent infection. This work provides mechanism by which HCMV modulates the host cell environment to its advantage.

## Introduction

Human cytomegalovirus (HCMV), a β–herpesvirus, can infect a wide range of cell types, including undifferentiated myeloid cells, which represent an established, natural site of latent infection (1, 2). This latent infection underpins a lifelong, persistent infection with HCMV, present in ∼50–75% of the human population (3). Although healthy individuals infected with HCMV are predominantly asymptomatic, lytic reactivation from latency causes significant health complications and often mortality in immune-deficient patients (4, 5). The threat of HCMV reactivation in immunocompromised patients is most acute in transplant recipients (6, 7). While lytic HCMV infection is reduced and often controlled with anti-viral therapies, these treatments target the lytic phase of infection, are problematically toxic, and are rendered less effective due to emerging drug resistant strains (8, 9). Thus, defining novel means to prevent viral reactivation will therefore have significant clinical benefits.

Latent infection is defined as the maintenance of viral genomes in the absence of infectious virion production, coupled with the ability to reactivate lytic replication to produce infectious virus, given the proper extracellular and/or environmental cues (10). As CMVs are highly host specific, HCMV latency studies are largely restricted to models utilizing human cells, including *ex vivo* primary hematopoietic cells, such as monocytes (11–13) and CD34^+^ hematopoietic progenitor cells (HPCs) (14–16), as well as *in vitro* model systems (17–23), such as Kasumi-3 cells (22) and embryonic stem cells (23). Recent advancements in humanized mouse model systems have begun to allow for *in vivo* HCMV studies (24–28). Defining the HCMV latent transcriptome represents a key area of continued research, and at present, the exact roles of many latency associated genes still remain unclear. However, genes such as *UL135*, *UL138*, *US28*, *LAcmvIL-10* (latency-associated HCMV IL-10 homolog), latency unique natural antigen (*LUNA*), and *UL144* play a role in maintaining HCMV latency and subsequent reactivation (29). HCMV also manipulates cellular factors to maintain latent infection, including cellular microRNAs (30), cell surface protein expression (31–34), and the cellular secretome (35). Chromatinization of the major immediate early promoter (MIEP) during latency contributes to securing extended viral transcriptional silencing and establishment of latency (36), yet how this occurs is still under investigation.

The HCMV *US28* gene is transcribed during both lytic and latent infection, during which the protein aids in securing latency in undifferentiated myeloid cells (37, 38). The *US28*-encoded protein (pUS28) is one of four G-protein coupled receptors (GPCRs) encoded by the HCMV genome (39) and is a potent signaling protein in the context of lytic infection (40). pUS28 is incorporated into the mature viral particle (37) and transcripts are expressed during both experimental (11, 15, 37, 38, 41–44) and natural latency (42, 45, 46). Although it is clear that pUS28 is necessary for HCMV latency (37, 38), whether this protein is required for the establishment and/or the maintenance of latency remains unknown. In this current study, we report that pUS28 functions to both establish and maintain viral latency. Additionally, our data reveal that pUS28 attenuates cellular fos (c-fos), a component of the AP-1 transcription complex that binds to the MIEP during lytic replication. Our data reveal that pUS28-mediated attenuation of c-fos activation results in a decrease in AP-1 binding to the MIEP. Together, our findings reveal a novel mechanism by which pUS28 coopts the host cell to silence MIEP-driven transcription and secure viral latency.

## Methods

### Cells and Viruses

Details on cells and their culture conditions, as well as details on the propagation of the TB40/E BAC-derived viruses are included in the SI Materials and Methods.

### Generation of stable NuFF-1 and THP-1 cell lines

For details on the generation of the pBABE-US28-3xF and pSLIK-US28-3xF constructs, as well as the US28-3xF expressing NuFF-1 cells, see SI Materials and Methods.

To generate the stable THP-1-pSLIK-hygro and THP-1-pSLIK-US28-3xF lines, pSLIK-hygro or pSLIK-US28-3xF were transfected into 293T cells using Fugene 6 Transfection Reagent (Promega), as detailed above. The clarified media was then concentrated 50X by ultracentrifugation at 82,700 × *g* at 4°C for 60 minutes (min), after which the pellet was resuspended in X-VIVO15 (Lonza). The concentrated pSLIK-hygro or pSLIK-US28-3xF lentiviral particles were then used to infect 5 × 10^5^ THP-1 cells per condition by centrifugal enhancement for 1 hour (h) at room temperature at 1000 × *g*. Successfully transduced cells were selected with 450 μg/ml hygromycin, after which cell debris was removed by cushioning cells onto Ficoll Paque™ PLUS (GE Healthcare Life Sciences) for 35 min at room temperature at 450 × *g* without the brake.

### RNA and Protein Analyses

For details on RNA and protein analyses, see SI Appendix, SI Materials and Methods.

### Infection of THP-1, Kasumi-3, and CD34^+^ cells

THP-1 cells were infected as described elsewhere (47). In brief, 1 × 10^6^ cells were infected at a multiplicity of 1 TCID_50_/cell by centrifugal enhancement at 450 × *g* for 35 min, without the brake, at room temperature. Infections were then returned to the culture conditions described in the SI Materials and Methods for an additional 75 min, after which the cells were washed with 1X PBS, replated, and incubated in X-VIVO15 media as indicated in the text. Doxycycline (DOX) induction of pUS28 expression, where stated, was performed 48 h prior to infection by adding 1 μg/ml DOX to 1 × 10^7^ cells in RPMI. After infection, transduced THP-1 cells were washed and replated in X-VIVO15 media with 1 μg/ml DOX. During the course of infection, transduced THP-1 cells were treated with DOX every two days after infection, and media was changed every 5 days (d), where cellular debris was removed by centrifuging cells onto Ficoll Paque™ PLUS (GE Healthcare Life Sciences) for 35 min at room temperature at 450 × *g* without the brake.

Kasumi-3 cells were infected as described previously (22, 30, 37). Briefly, cells were serum starved in X-VIVO15 48 h prior to infection, and then infected at a multiplicity of 1.0 TCID_50_/cell by centrifugal enhancement (1000 × *g*, 35 min, room temperature, with no brake) in X-VIVO15. The following day, cells were treated with trypsin to remove any virus that had not entered the cell, and then cushioned onto Ficoll-Pacque (GE Healthcare) to remove residual virus and debris. Infected cells were washed with 1X PBS, replated in X-VIVO15, and harvested as indicated in the text.

Isolation of CD34^+^ HPCs is described in detail elsewhere (48). Immediately following isolation, CD34^+^ HPCs were infected at a multiplicity of 2.0 TCID_50_/cell, as previously described (22, 30, 37, 48), in infection media consisting of IMDM (Corning) supplemented with 10% BIT9500 serum substitute (Stem Cell Technologies), 2 mM L-glutamine, 20 ng/ml low-density lipoproteins, and 50 μM 2-mercaptoethanol. The next day, cultures were washed three times in 1X PBS, and replated in 0.4μm pore transwells over irradiated stromal cells in hLTCM, as described previously. Following 12 d post-infection (dpi), cells were washed three times in 1X PBS and harvested for RT-qPCR analyses and extreme limiting dilution assays (ELDA).

### Assaying the Release of Infectious Virus by ELDA

Infected Kasumi-3 cells or CD34^+^ HPCs were co-cultured with naïve NuFF-1 cells. Briefly, infected cells were serially diluted 2-fold onto naïve NuFF-1 cells in X-VIVO15 for 14 d. The number of mCherry-positive wells for each dilution was counted and virus release measured using ELDA software (http://bioinf.wehi.edu.au/software/elda/index.html).

### Chromatin Immunoprecipitation

Cells were infected for 24 h with WT or US28Δ, fixed in 1% formaldehyde for 15 min at room temperature, and then quenched with 125 mM glycine. Cells were lysed in IP buffer (150 mM NaCl, 50 mM Tris-HCl pH 7.5, 5 mM EDTA, 0.5% Igepal-CA630, and 1% Triton-X-100) and debris removed by centrifugation. DNA was sheared to 0.3-1 kB fragments with a MiSonix Sonicator 3000 (20% output, 0.5 s on/off, 1 min) and aliquots stored as input controls. DNA associated with AP-1 was immunoprecipitated with either normal rabbit serum (Cell Signaling) or anti-fos (ChIP grade; Cell Signaling, diluted 1:100) using protein A agarose (Millipore Sigma). DNA was eluted by boiling and was followed by proteinase K treatment. DNA from disrupted nucleosomes was precipitated and used in PCR with primers directed at the MIEP or the UL69 non-promoter region (49) (Supplemental Appendix, Table S1).

## Results

### Exogenous pUS28 expression rescues the lytic growth phenotype in US28Δ-infected myeloid progenitor cells

To begin to understand the role of *US28* during establishment and/or maintenance of latency, we transduced THP-1 cells with a lentiviral construct (pSLIK-US28-3xF) that allows for inducible expression of pUS28 fused to a C-terminal triple FLAG epitope tag. We first assessed the efficiency of pUS28 expression in this system by treating THP-1-pSLIK-US28-3xF or THP-1-pSLIK-hygro cells with or without DOX and then harvested the cells over 24 h. We detected pUS28 expression within 1 h of induction, with the most robust expression observed 24 h post-treatment. Importantly, the expression of pUS28 is tightly regulated in this inducible system, as we did not observe pUS28 in the absence of DOX (Fig 1A). Additionally, to understand pUS28 degradation kinetics in this system, we treated THP-1-pSLIK-US28-3xF cells with DOX for 24 h (day 0), then washed the cells to remove the DOX, and harvested these cells every 2 d for 6 d to monitor pUS28 levels. Our findings indicate that pUS28 is significantly degraded 2 d after the removal of DOX and is undetectable by d 4 (Fig 1B).

**Figure 1.**
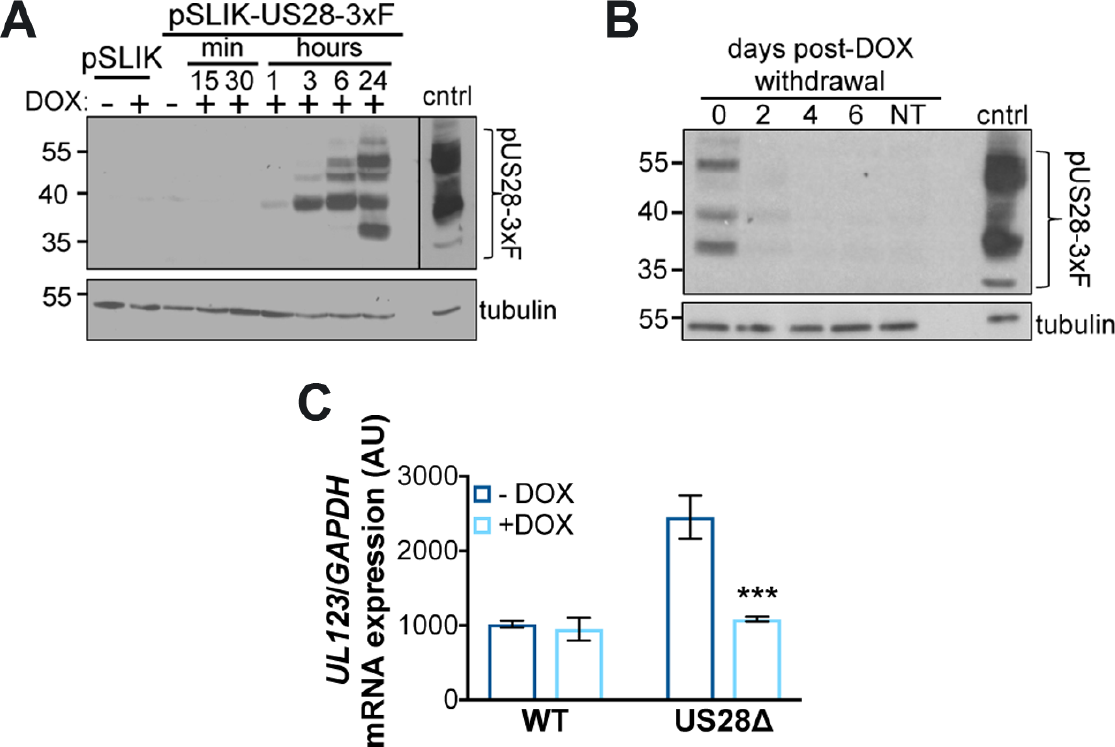
Continual pUS28 expression complements the lytic-like phenotype following US28Δ infection. (**A**) THP-1 cells were transduced with THP-1-pSLIK-hygro (pSLIK) or THP-1-pSLIK-US28-3xF (pSLIK-US28-3xF) and treated with (+) or without (−) DOX (1 μg/ml) to induce pUS28 expression. Cells were harvested at the indicated time points post-treatment. pUS28 expression was detected by immunoblot using the FLAG epitope tag. (**B**) THP-1-pSLIK-US28-3xF cells were treated with DOX for 24 h and harvested to confirm pUS28 expression (0 d). Remaining cells were washed in PBS and cultured in the absence of DOX for the duration of the experiment. Cell samples were taken at the indicated days post-treatment and all samples were then immunoblotted for pUS28 using the epitope tag. (**A, B**) As a control (cntrl), NuFF-1 cells were infected with US28-3xF (moi = 0.5) and cell lysates were harvested at 96 hpi. Tubulin is shown as a loading control. Note that due to the intensity of this control lysate in (**A**), a shorter exposure is shown, as denoted by the black line. (**C**) THP-1-pSLIK-US28-3xF cells were treated with (+; dark blue) or without (-; light blue) DOX for 24 h and then infected with WT or US28Δ (moi = 1.0). DOX was replenished every 48 h and the cells were harvested at 6 dpi for *UL123* expression by RT-qPCR. Samples were normalized to *GAPDH*, and each sample was analyzed in triplicate. Errors bars indicate standard deviation; *** *p* < 0.001.

We have previously shown that the deletion of the *US28* ORF results in a lytic rather than latent infection of both Kasumi-3 and CD34^+^ HPCs (37). Thus, to determine if pUS28 provided *in trans* could complement this phenotype, we infected THP-1-pSLIK-US28-3xF cells in the presence or absence of DOX with either TB40/E*mCherry* (WT) or TB40/E*mCherry*-US28Δ (US28Δ). We collected the cells 7 dpi and assessed viral lytic gene expression (*UL123*), as this serves as an indicator of MIEP activity level, and thus is a proxy for distinguishing a latent versus a lytic phenotype. Indeed, we found that DOX-induced THP-1-pSLIK-US28-3xF cells infected with US28Δ resulted in similar levels of *UL123* transcript levels when compared to WT infections (Fig 1C). This finding suggests that pUS28 expression helps to maintain the suppression of viral lytic gene expression.

### pUS28 expression at the time of infection fails to maintain MIEP repression

Our previous findings revealed that pUS28 is incorporated into the mature viral particle and is delivered to Kasumi-3 cells upon infection (37). Thus, we hypothesized that virion-delivered pUS28 may function early post-infection to aid in establishing a latent infection in myeloid cells. To test whether virion-delivered pUS28 could establish and/or maintain latent infections, we generated a pUS28 complemented recombinant virus, which incorporates pUS28 into the virion, but lacks the *US28* ORF (TB40/E*mCherry*-US28*comp*; US28*comp*) by growing US28Δ on a pUS28 expressing fibroblast cell line (Fig S1A, S1B). We confirmed that US28*comp* virus incorporates pUS28 into the virion (Fig S1C) and that it does not express *de novo US28* mRNA after infection of NuFF-1 cells (Fig S1D). We noticed that pUS28 expression is marginally lower in US28*comp* virus in comparison to US28-3xF virion incorporated pUS28 (Fig S1C), which could be due to pUS28 expression levels in the complementing cell line. Nonetheless, these data show that US28*comp* incorporates pUS28 into the mature viral particle, but unable to produce *de novo* pUS28 upon subsequent infection.

Using this novel, complemented virus, we asked if virion-delivered pUS28 was capable of suppressing lytic gene expression in conjunction with our THP-1 cells overexpressing US28. To this end, we treated THP-pSLIK-US28-3xF with or without DOX and then infected them with WT, US28Δ, or US28*comp* and either maintained DOX treatment for 12 d, or removed DOX treatment after infection. We found that low *UL123* expression correlated with the maintenance of pUS28 expression, either by DOX treatment or by infection with WT virus. However, in cultures where pUS28 was not continually expressed for the duration of the experiment, *UL123* expression was significantly higher (Fig 2A). This demonstrates that pUS28 expression must be sustained in order to maintain repression of MIEP-driven transcription during HCMV infection of monocytic cells. Correlating with these findings, *UL99* expression followed a similar pattern to *UL123* (Fig 2B). We also assessed the ratio of *UL138*/*UL123* mRNA expression (22, 37) to discern those cultures that favor a latent transcriptional profile (high *UL138*/*UL123* ratio) from those that favor a lytic transcriptional profile (low *UL138*/*UL123* ratio. The *UL138*/*UL123* ratios for the US28Δ- and US28*comp*-infected cells that were cultured in the absence of exogenous pUS28 expression were significantly lower than the same cultures that received DOX after infection, as well as the WT-infected cells, regardless of treatment (Fig 2C). Together, these findings argue that virion-delivered pUS28, or pUS28 expressed only at the time of infection, fails to maintain the suppression of lytic gene transcription.

**Figure 2.**
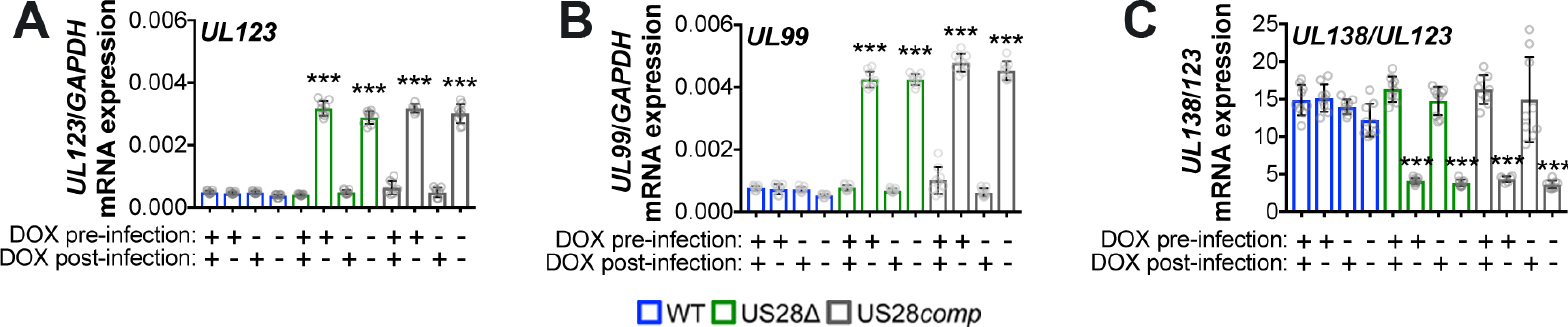
Virion-delivered pUS28 fails to maintain suppression of lytic gene transcription. THP-1-pSLIK-US28-3xF were treated with (+) or without (−) DOX (1 μg/ml) to induce pUS28 expression (pre-infection), and then infected with WT (blue), US28Δ (green), or US28*comp* (gray) (moi = 1.0). Infected cells were washed 2 hpi and cultured in the presence (+) or absence (−) of DOX, as indicated (post-infection), where cultures receiving post-infection DOX were replenished every 48 h. Cells were harvested 12 dpi and (**A**) *UL123*, (**B**) *UL99*, and (**C**) the ratio of *UL138*/*UL123* expression were measured by RT-qPCR. Samples were normalized to *GAPDH* and analyzed in triplicate. Errors bars indicate standard deviation and statistical significance was calculated using one-way ANOVA and Dunnett’s post-hoc analysis; \*\*\**p* < 0.01.

### pUS28 expression results in repression of MIEP-driven transcription

Based on our above findings, we hypothesized that pUS28 functions to repress the MIEP, both at early time points after infection to establish latency as well as throughout infection to maintain latency. To test this, we pre-treated THP-1-pSLIK-US28-3xF cells with or without DOX for 48 h to induce pUS28 expression. We then infected these cells with either WT or US28Δ (multiplicity of infection [moi] = 1.0) and cultured the cells for 7 d, with or without continued DOX treatment. Cells were then washed and DOX treatment was either maintained or reversed for each culture, as indicated. We observed that continual pUS28 expression throughout infection maintained repression of the MIEP, as measured by *UL123* for WT-versus US28Δ-infected cells cultured in the presence of DOX (Fig 3A). Secondly, removal of DOX-induced pUS28 expression after 7 d of treatment led to a de-repression of *UL123* transcription by the end of the assay at day 14 (Fig 3A), thus confirming the requirement for pUS28 in maintaining repression of MIEP-driven transcription. Finally, we observed that introduction of DOX-induced pUS28 expression for the last 7 d in US28Δ-infected cells, initially cultured in the absence of DOX, lead to an intermediate phenotype (Fig 3A), suggesting that introduction of DOX-induced pUS28 expression later in infection can repress MIEP-driven transcription.

**Figure 3.**
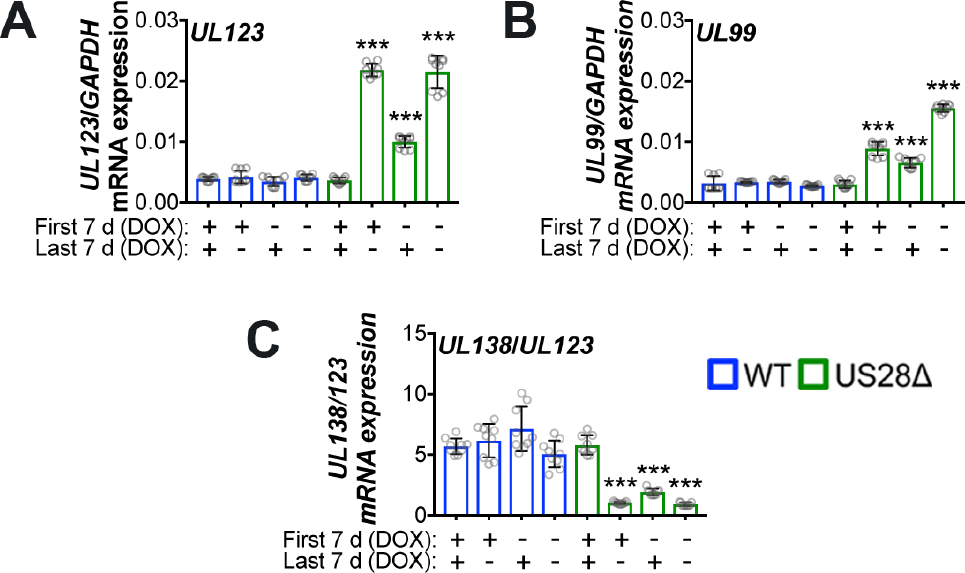
Sustained pUS28 expression suppresses lytic gene expression. THP-1-pSLIK-US28-3xF cells were treated with (+) or without (−) DOX (1 μg/ml) for 48 h to induce pUS28 expression and then infected with WT (blue) or US28Δ (green) (moi = 1.0). Cells were cultured for 7 d, during which cells were maintained under their original treatment conditions (First 7 d). At 7 dpi, cells were washed and treated with (+) or without (−) DOX (Last 7 d). Total RNA was harvested and (**A**) *UL123*, (**B**) *UL99*, and (**C**) the ratio of *UL138*/*UL123* expression were measured by RT-qPCR. Samples were normalized to *GAPDH* and assessed in triplicate. Errors bars indicate standard deviation and significance calculated using one-way ANOVA and Dunnett’s post-hoc analysis; \*\*\**p* < 0.001.

We also assessed the expression of *UL99*, an HCMV gene associated with late lytic infection, and found this gene is also repressed in response to pUS28 expression (Fig 3B). Interestingly, when pUS28 expression is induced by DOX treatment for the first 7 d of infection followed by DOX withdrawal, *UL99* mRNA expression increases, although it does not reach the levels we observed in the absence of DOX-induced pUS28 expression. In agreement with *UL123,* introduction of DOX-induced pUS28 expression only for the last 7 d of infection, leads to a reduction in *UL99* mRNA expression (Fig 3B). Taken together, these data suggest that the loss of pUS28 expression results in the de-repression of MIEP-driven *UL123* expression, leading to induction of the lytic lifecycle, including expression of the late gene products. Furthermore, assessing the ratio of *UL138*/*UL123* mRNA expression (22, 37) confirmed the correlation between those cultures that favor a latent transcriptional profile (high *UL138*/*UL123* ratio) versus those that favor a lytic transcriptional profile (low *UL138*/*UL123* ratio) (Fig 3C). Additionally, pUS28 fails to repress lytic gene transcription in differentiated THP-1 cells in which the virus lytically replicates (50) (Fig S2 and S3), suggesting that this effect is specific to undifferentiated cells. Together, these data suggest that pUS28 suppresses lytic transcription in cells that support latency.

### pUS28 represses the MIEP transcription at times consistent with the establishment of latent infection in Kasumi-3 cells

Having observed that ectopic expression of pUS28 represses *UL123* and *UL99* expression in THP-1 cells, we asked whether pUS28 plays a role in suppressing lytic gene transcription during latent infection of Kasumi-3 cells, a culture system that affords us the ability to assess both functional latency and reactivation (22). We asked if virion-delivered pUS28 was capable of suppressing lytic gene expression in the context of infection and whether pUS28 contributes to the establishment of latency. We infected Kasumi-3 cells with WT, US28Δ, or US28*comp* (moi = 1.0) and measured expression of *UL123*, *UL99*, and *UL138*/*UL123* over 12 d. Firstly, we observed a burst in *UL123* and *UL99* mRNA expression following WT infection, which peaked 3 dpi (Fig 4A, 4B), as previously observed by others (11, 15, 51). *UL123* and *UL99* gene expression was lower in Kasumi-3 cells infected with WT or US28*comp* viruses at this time point (Fig 4A, 4B), suggesting that virion delivered pUS28 has a repressive effect on lytic transcripts at early time points post-infection.

**Figure 4.**
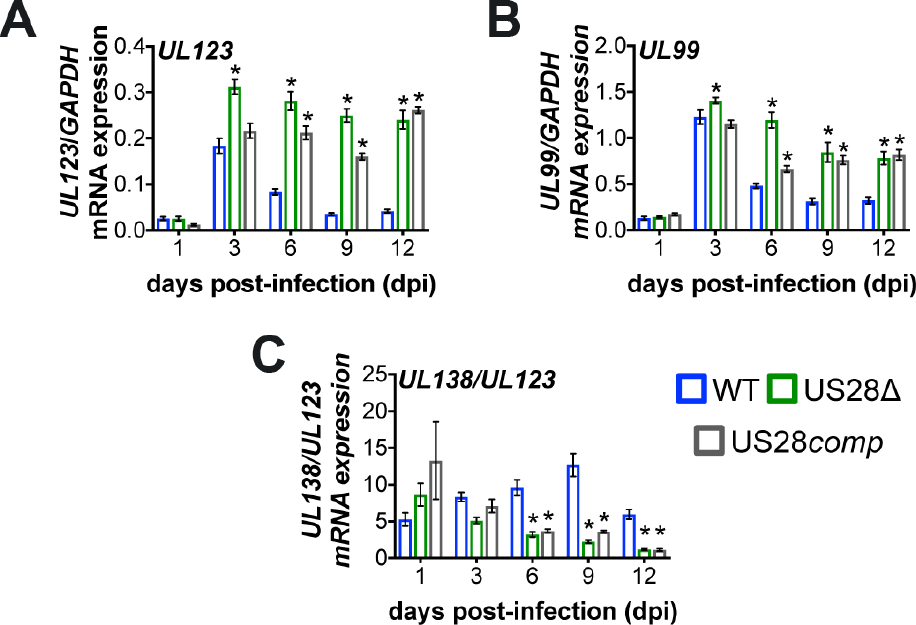
pUS28 represses *UL123* and *UL99* expression in infected Kasumi-3 cells at early times of latent infection. Kasumi-3 cells were infected with WT (blue), US28Δ (green), or US28*comp* (gray) (moi = 1.0) and sorted for mCherry positive cells at 1 dpi. Cells were then harvested at the indicated dpi. (**A**) *UL123*, (**B**) *UL99*, and (**C**) the ratio of *UL138*/*UL123* expression were measured by RT-qPCR. Samples were normalized to *GAPDH* and analyzed in triplicate. Errors bars indicate standard deviation, and statistical significance was calculated using two-way ANOVA and Dunnett’s post-hoc analysis relative to WT virus at each time point; * *p* < 0.05.

Subsequent to this initial burst of mRNA expression, transcription of these genes decreased in the WT-infected cells compared to US28Δ- or US28*comp*-infected Kasumi-3 cells. Additionally, *UL123* expression stabilized in US28Δ-infected cells, at an expression level that is higher than that of cells infected with WT (Fig 4A), suggesting that pUS28 represses lytic gene transcription at early times post-infection of Kasumi-3 cells, at times consistent with the establishment of latency. Additionally, these data suggest that continued pUS28 expression represses lytic gene transcription to contribute to maintaining latency. Confirming this hypothesis, pUS28 that is delivered by US28*comp* virus is capable of repressing *UL123* and *UL99* expression at early times post-infection (Fig 4A, WT vs. *US28comp* at 3 dpi), but failed to repress these lytic gene products from 6 dpi onwards (Fig 4A and 4B), suggesting that repression is diminished after virion-delivered pUS28 is degraded. Finally, the ratio of *UL138*/*UL123* significantly lower in US28Δ- or US28*comp*-infected compared to WT-infected Kasumi-3 cells from 6 dpi through the duration of the experiment, supporting the findings that these cultures display a more lytic phenotype (Fig 4C). Overall, these data suggest that pUS28 represses MIEP-driven transcription, which helps to both establish and maintain latency.

### pUS28 repression of the MIEP reduces virus production

Our data so far reveal that pUS28 suppresses lytic gene transcription in cells that support viral latency. To determine if pUS28 aids in establishing and maintaining latent infection, we took advantage of the *in vitro* Kasumi-3 cell and *ex vivo* primary cord blood-derived CD34^+^ HPC latency models. First, we infected Kasumi-3 cells with WT, US28Δ, or US28*comp* (moi = 1.0) and measured the production of infectious particles by extreme limiting dilution assay (ELDA) by co-culture with fibroblasts, sampling every 3 dpi for 12 d. Our results indicate that US28*comp*-infected Kasumi-3 cells display a similar phenotype to WT-infected Kasumi-3 cells until 6 dpi, after which the US28*comp* infections reveal an intermediate phenotype, releasing significantly more virus than WT infections, indicating a failure to maintain latency (Fig 5A). To confirm these findings in a cell type that represents a natural site of HCMV latency, we infected cord blood-derived CD34^+^ HPCs with WT, US28Δ, or US28*comp* (moi = 2.0) and cultured the cells for 12 d under conditions that promote latency. We then harvested the cells for ELDA analysis on naïve fibroblasts, as above. We found that US28*comp*-infected CD34^+^ HPCs produce infectious virions to similar levels as we observe in the US28Δ-infected CD34^+^ HPCs (Fig 5B). Consistent with these data, viral lytic gene expression in Kasumi-3 cells and CD34^+^ HPCs at 12 dpi revealed increased *UL123* and *UL99* in US28Δ- or US28*comp*-infected cultures, compared to cells infected with WT (Fig S4). These data, suggest that virion-delivered pUS28 contributes to establishing HCMV latency, however it is not sufficient to maintain long-term latent infections, and therefore, sustained pUS28 expression is required to suppress viral replication.

**Figure 5.**
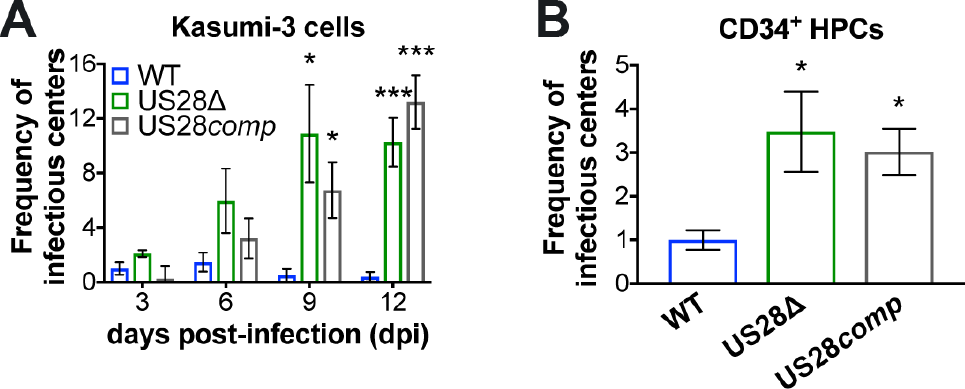
Sustained pUS28 expression is required to maintain viral latency. (**A**) Kasumi-3 cells (moi = 1.0) or (**B**) CD34^+^ HPCs (moi = 2.0) were infected with WT (blue), US28Δ (green), or US28*comp* (gray). (**A, B**) Cells were collected at the indicated time points and reactivation events were determined by ELDA. Data in (**A**) is presented as fold-change in virus release relative to WT virus on day 3. Data in (**B**) is presented as fold change in virus release relative to WT virus. In all cases, error bars represent standard deviations of three biological replicates. Statistical significance was calculated relative to WT at the same time point using (**A**) two-way ANOVA analysis and Dunnett’s post-hoc analysis in or (**B**) one-way ANOVA. * *p* < 0.05, *** *p* < 0.001.

### pUS28 expression is sufficient to attenuate c-fos expression and signaling

Our previous findings revealed that pUS28 is a potent signaling molecule during the context of infection (40). Thus, we hypothesized that pUS28’s signaling capabilities contribute to its function during latency. However, our previous work also noted that pUS28-mediated signaling is cell type specific (40), making it difficult to extrapolate such findings to hematopoietic cells. To begin to understand the pUS28-modulated pathways and the mechanism underlying pUS28’s contribution to establishing and maintaining latency, we performed a PCR array analysis on THP-pSLIK-US28-3xF, compared to control cells, treated with DOX for 24 h to induce pUS28 expression. This analysis revealed a subset of statistically significant up-regulated and down-regulated cellular genes (Fig S5). The most significantly up-regulated genes in the Pathway Analysis were *CHUK* (conserved helix-loop-helix ubiquitous kinase; IKKα), *HRAS*, *JAK2* (janus kinase 2), *PIK3CB* (phosphatidylinositol-4,5-bisphosphate 3-kinase catalytic subunit beta), and *STAT1* (signal transducer and activator of transcription 1), while *c*-*fos*, *PIK3CA* (PIK3C alpha), *src*, and *STAT5A* were the most significant down-regulated genes in response to pUS28 expression (Fig S5).

We focused on *c-fos* since activation of c-fos results in the heterodimerization of c-fos and c-jun to form the activator protein 1 (AP-1) complex, which binds to the MIEP, thereby aiding in its transcriptional activation (52). Thus, we hypothesized that pUS28-mediated attenuation of *c-fos* transcription reduces AP-1 complex formation, thus leading to the promotion of lytic transcription. We confirmed that *c-fos* expression is reduced in transduced THP-1 cells upon induction of pUS28 by RT-qPCR (Fig 6A). We also assessed *c-jun* expression, which is significantly downregulated during latent infection of CD34^+^ HPCs and Kasumi-3 cells during HCMV infection (53). Interestingly, we found pUS28 expression did not significantly impact *c-jun* (Fig 6A), suggesting that pUS28 specifically targets *c-fos* and not both AP-1 components, and that another as of yet unknown viral factor likely targets *c-jun*. We found that pUS28 also reduces c-fos protein expression and significantly decreases c-fos phosphorylation (Fig 6B). Interestingly, pUS28 expression does not affect c-fos or c-jun expression in differentiated THP-1 cells (Fig S7A), suggesting that pUS28-mediated regulation of c-fos is specific to cells that support HCMV latency.

**Figure 6.**
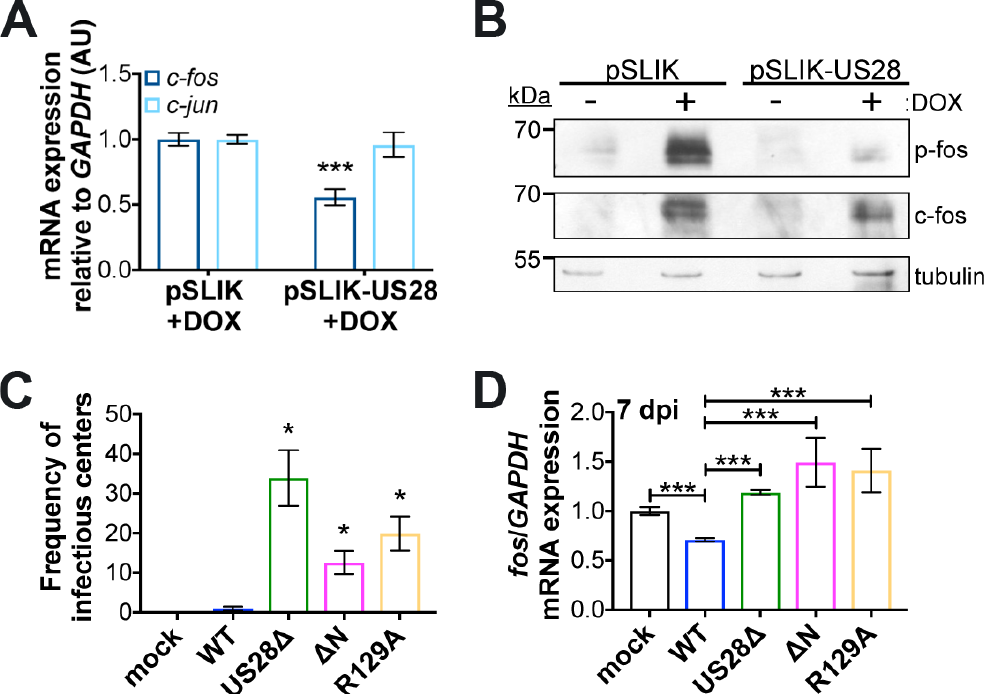
US28 signaling attenuates both *c-fos* expression and activation. (**A**) THP-1-pSLIK-hygro (pSLIK) and THP-1-pSLIK-US28-3xF (pSLIK-US28) were treated with DOX and cells were harvested 24 h post-treatment. *c-fos* (dark blue) and *c-jun* (light blue) expression were measured relative to *GAPDH*. (**B**) Cells from (**A**) were treated with (+) or without (−) DOX and cells were harvested 24 h post-treatment. Phosphorylated fos (p-fos), total fos (c-fos), and tubulin were detected by immunoblot. (**C, D**) Kasumi-3 cells were infected with the indicated viruses (moi = 1.0) and harvested at 7 dpi. (**C**) The frequency of infectious virus from each latently infected culture was determined by ELDA. Data is presented as fold-change in virus release relative to WT. (**D**) *c-fos* expression was measured relative to *GAPDH*. (**A, C, D**) Each sample was analyzed in triplicate. Error bars indicate standard deviation. Statistical significance was calculated using one-way ANOVA and Dunnett’s post-hoc analysis; \**p* < 0.05, \*\**p* < 0.01, \*\*\**p* < 0.001.

### Latent expression of pUS28 signals to attenuate c-*fos* expression

To determine if pUS28-mediated signaling attenuates c-fos in the context of infection, we took advantage of two additional US28 viral recombinants that alter the ability of US28 to signal. US28-R129A has a single point mutation at amino acid position 129 in the canonical “DRY” motif (37). Mutation of this motif from DRY to DAY prevents G proteins from coupling to this domain, rendering this mutant ‘G-protein signaling-dead’ (54). The US28ΔN recombinant lacks N-terminal amino acids 2-16 (37) and is unable to interact with US28 ligands, though this mutant can still signal constitutively (54). We generated each of these signaling mutants in the TB40/E*mCherry*-US28-3xF background, and confirmed pUS28 expression during fibroblast infection with each recombinant by immunoblot (Fig S6A). While there was a slight growth advantage with the US28-R129A mutant at 8 dpi, there was no significant difference between either mutant and wild type by 15 dpi (Fig S6B). Next, we infected Kasumi-3 cells under conditions that promote latency with WT, US28Δ, US28ΔN, or US28-R129A and measured the release of infectious virus by ELDA. WT-infected cells established and maintained a latent infection, while US28Δ-, US28-R129A-, and US28ΔN-infected cells all resulted in significant lytic replication (Fig 6C), suggesting that chemokine binding and agonist-dependent signaling is required. To determine if pUS28-mediated signaling impacts *c-fos* expression, we latently infected Kasumi-3 cells with WT, US28Δ, US28ΔN, or US28-R129A and measured *c-fos* expression 2 and 7 dpi. WT virus attenuated *c-fos* expression as early as 2 dpi, whereas *c-fos* expression increased in US28Δ-, US28-R129A-, and US28ΔN-infected cells (Fig S7B). These findings were amplified by 7 dpi, confirming that continued expression of pUS28 attenuates *c-fos* and pUS28-mediated signaling is important for *c-fos* regulation (Fig 6D).

Given our observations that 1) pUS28 attenuates *UL123* expression, 2) pUS28 attenuates *c-fos* expression and activation, and 3) there are AP-1 binding sites in the MIEP (55), we hypothesized that pUS28 expression in cells that support latency results in a reduction of AP-1 binding to the MIEP. To this end, we infected Kasumi-3 cells with WT or US28Δ and performed chromatin immunoprecipitation (ChIP) for AP-1 at the MIEP at 2 and 7 dpi using an antibody directed at c-fos. Our data reveal an increase in AP-1 bound to the MIEP in US28Δ-infected cells as early as 2 dpi (Fig S8), which is maintained through 7 dpi (Fig 7A). Finally, we reasoned that if pUS28-mediated attenuation of c-fos signaling suppresses MIEP activation, then treatment of US28Δ-infected cells with a c-fos inhibitor should reduce MIEP activity. Infection of Kasumi-3 cells, with either WT or US28Δ in the presence of a c-fos inhibitor (T-5224) for 7 d showed reduced *UL123* expression in US28Δ-infected cultures (Fig 7B), similar to levels we observed in the presence of pUS28 exogenous expression (Fig 6A). Additionally, this reduction in *UL123* expression in the US28Δ-infected Kasumi-3 cells results in a decrease in the production of infectious virus compared to its untreated counterpart (Fig 7C). Together, these data show that WT virus significantly reduces AP-1-mediated activation of the MIEP by pUS28-directed attenuation of *c-fos*, thereby contributing to the establishment and maintenance of latency in cells that support HCMV latent infection.

**Figure 7.**
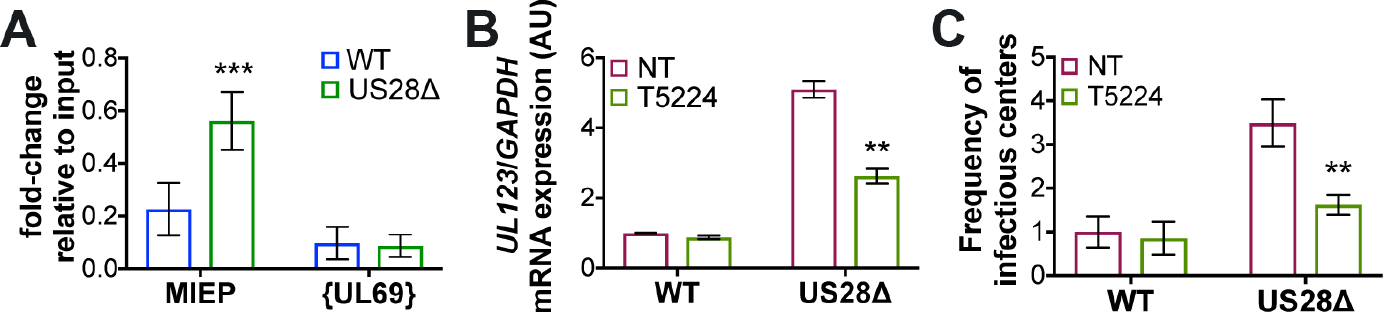
pUS28 decreases c-fos binding at the MIEP, leading to transcriptional repression of IE transcripts. (**A**) Kasumi-3 cells were infected (moi = 1.0) with WT (blue) or US28Δ (green). Cells were collected 7 dpi and the AP-1 complex was immunoprecipitated using an anti-c-fos antibody. Co-precipitated MIEP was quantified by qPCR, and data is shown as fold-change relative to input. The UL69 non-promoter region is shown as a control. (**B,C**) Kasumi-3 cells were infected as in (**A**) in the absence (red; NT) or presence (green) of the fos inhibitor, T5224 (10 nM), and cells were harvested 7 dpi. (**B**) *UL123* expression was measured and normalized to *GAPDH*. (**C**) The frequency of infectious virus from each latently infected culture was determined by ELDA. Data is presented as fold-change in virus release relative to vehicle-treated WT. Each sample was analyzed in triplicate. Errors bars indicate standard deviation and statistical significance was measured using Welch’s T-test; * *p* < 0.05, \*\**p* < 0.01.

## Discussion

Deciphering the molecular underpinnings of the establishment and maintenance of HCMV latency is an on-going area of intense research. Viral infection of hematopoietic cells results in what is likely a highly coordinated hijacking of the host cell environment, whereby multifaceted processes dictate the infection outcome. The primary finding in this study is a requirement for pUS28 during both the establishment and maintenance phases of HCMV latency, dictated, at least in part, by its manipulation of host encoded c-fos. pUS28 also influences AP-1 binding to the MIEP, a cellular transcription factor that is comprised of c-fos and c-jun subunits. Finally, pUS28-mediated attenuation of c-fos directly impacts MIEP-driven transcription. Our results reveal a novel mechanism by which pUS28 functions to suppress MIEP-driven transcription to aid in the establishment and maintenance of HCMV latency.

Viral establishment and maintenance of latency likely requires a multitude of factors, and pUS28 is a key piece of the biological puzzle that balances latency and reactivation. Suppression of the very strong lytic promoter, the MIEP, is a major determinant for a successful latent infection, and pUS28 is certainly not the only viral factor involved in this mechanism. Epigenetic silencing of the MIEP is well-established and additional viral factors, including non-coding RNAs and proteins, clearly influence cellular factors that regulate this viral promoter during infection of cells that support latency (56). While host cell proteins like Ying Yang-1 (YY-1) and TRIM28/Kap-1 have been shown to impact MIEP transcription and viral latency, the upstream factors that lead to their regulation remain elusive, though it is attractive to speculate that viral-induced signaling, by way of pUS28, may be involved.

pUS28-mediated attenuation of the AP-1 transcription factor complex is specific to c-fos, as c-jun was unaffected. In a recent collaboration, we found that *c-jun* expression is significantly down-regulated during HCMV latent infection of both Kasumi-3 cells and primary CD34^+^ HPCs (53), although we show herein that pUS28 expression did not significantly alter *c-jun* expression. Together, these data suggest another viral factor regulates *c-jun* transcript levels during latency. HCMV has therefore devised multiple mechanisms to regulate the AP-1 complex, suggesting this transcription factor is important to regulating latency. In addition to preventing AP-1 binding at the MIEP, it will be interesting to further dissect the biological consequences of pUS28-mediated c-fos regulation. More global, unbiased approaches, such as RNAseq and whole cell proteomics may reveal additional host cell factors that are dysregulated via the pUS28:c-fos signaling axis. Indeed, these downstream events may provide an additional mechanism by which HCMV regulates the host cell during latency.

Our data also reveal that pUS28-mediated signaling is important for maintaining latency, through the regulation of *c-fos* in both a G-protein coupling- and ligand binding-dependent fashion. The data we present herein agree with previous findings, which showed that US28-R129A (G-protein coupling domain point mutant) expression in THP-1 cells was unable to restore lytic gene suppression after infection with HCMV Titan-US28Δ virus (38). However, the role of ligand-mediated pUS28 signaling still remains unclear. Overexpression of the US28-Y16F ligand binding mutant in THP-1 cells suppresses lytic gene expression after infection with HCMV Titan-US28Δ virus (38). It is plausible that we have yet to identify a complete list of US28 ligands, thus perhaps the US28-Y16F mutant retains some ligand responsiveness, while the US28Δ mutant is completely devoid of such interactions.

Although US28 is linked to a variety of signaling pathways during lytic infection of different cell types (54), this is the first report that directly links pUS28 to c-fos signaling. Consistent with our findings, pUS28 expression in undifferentiated THP-1 cells attenuates MAP kinase signaling (38), a pathway that activates c-fos (57). Conversely, pUS28 activates this same pathway in differentiated THP-1 cells (38, 58, 59), which support lytic rather than latent infection. In line with this observation, we confirmed that pUS28’s repressive effect on the MIEP, via c-fos attenuation, is specific to undifferentiated myeloid cells that support latency, and does not occur in differentiated myeloid cells that support lytic infection. Whether MAP kinase signaling via pUS28 contributes to c-fos attenuation during the context of latent infection remains unknown, though it is attractive to hypothesize that pUS28’s impact on MAP kinase results in the downstream effects on c-fos expression we have observed. Furthermore, while it is possible that the alterations in the cellular environments between these different differentiation states of myeloid cells might explain the variations in US28 signaling activity, such conclusions remain tenuous.

How else might US28 repress the MIEP in myeloid progenitor cells? Given that pUS28 modulates a variety of signaling pathways during lytic infection (54), it is likely that future investigations will reveal additional pUS28-mediated cellular signaling pathways, some of which may aid in suppressing robust lytic gene transcription. In addition to MAP kinase signaling in THP-1s (38), US28 also activates signal transducer and activator of transcription 3-inducible nitric oxide synthase (STAT3-iNOS) in CD34+ cells (60) and PLC-β in THP-1s (61). How these pathways, along with others, might affect transcriptional suppression of the virus during latency in undifferentiated myeloid cells remains elusive.

Our findings described herein detail one specific mechanism by which pUS28 modulates the host cell environment to influence HCMV latency. Further work focused on understanding additional pUS28 activities during latent infection is essential to developing therapeutic targets aimed at preventing viral reactivation.

## Acknowledgments

This work was supported by National Institutes of Health grant AI119415 (to C.M.O’C.), American Heart Association Scientist Development Grant 15SDG23000029 (to C.M.O’C.), and University at Buffalo – SUNY and Cleveland Clinic funding (to C.M.O’C.).

